# Visible light optical coherence tomography angiography (vis-OCTA) and local microvascular retinal oximetry in human retina

**DOI:** 10.1101/2020.04.17.047514

**Authors:** Weiye Song, Wenjun Shao, Wei Yi, Rongrong Liu, Manishi Desai, Steven Ness, Ji Yi

## Abstract

We report herein the first visible light optical coherence tomography angiography (vis-OCTA) for human retinal imaging. Compared to the existing vis-OCT systems, we devised a spectrometer with a narrower bandwidth to increase the spectral power density for OCTA imaging, while retaining the major spectral contrast in the blood. We achieved a 100 kHz A-line rate, the fastest acquisition speed reported so far for human retinal vis-OCT. We rigorously optimized the imaging protocol such that a single acquisition takes <6 seconds with a field of view (FOV) of 3×7.8 mm^2^. The angiography enables accurate localization of microvasculature down to the capillary level and thus enables oximetry at vessels < 100 μm in diameter. We demonstrated microvascular hemoglobin oxygen saturation (sO_2_) at the feeding and draining vessels at the perifoveal region. The longitudinal repeatability was assessed by <5% coefficient of variation (CV). The unique capabilities of our vis-OCTA system may allow studies on the role of microvascular oxygen in various retinal pathologies.

## Introduction

Visible light optical coherence tomography (vis-OCT) [1–11] is a new derivation of OCT that uses visible light laser illumination instead of a conventional near-infrared (NIR) light [12]. One advantage of vis-OCT is its spatio-spectral analysis within the microvasculature for label-free oximetry (*i.e.* measuring hemoglobin oxygen saturation, sO_2_) [4,13–19]. Compared to 2D hyperspectral imaging modalities [20,21], vis-OCT’s precise 3D localization of blood vessels excludes a myriad of confounding factors from other tissue depths, enabling accurate and reliable oximetry measurements.

The first demonstration of vis-OCT dates back to 2002 [22], but the first *in vivo* vis-OCT retinal imaging was not reported until 2013 [4]. On the other hand, the development of vis-OCT oximetry has made significant progress in the past 10 years. The feasibility of using visible light for oximetry was first demonstrated *in vitro* in 2010 [23,24], and *in vivo* in 2011 [6]. Subsequently, the first demonstration of vis-OCT retinal oximetry was reported in 2013 in a rat retina, showing the sO_2_ calculations in the major arterioles and venules immediately exiting and entering the optic nerve head (ONH) [4]. When combined with the Doppler OCT measurements of blood flow, the oxygen metabolic rate from the inner retinal circulation can be quantified to access the global inner retinal functions and study the dynamic interaction between the retinal and choroidal circulations [3,15,25,26]. Later, human retinal imaging by vis-OCT was demonstrated in 2015 [27] and human retinal oximetry by vis-OCT was reported in 2017 [13,17].

So far, most reports on vis-OCT human retinal oximetry describe vessels in close proximity with the ONH with vessel diameters above 100 µm [13,17]. The arteriovenous difference in those major vessels provides a global assessment of inner retinal metabolic activity but is less informative for localized metabolic alterations. For example, primary open-angle glaucoma (POAG) frequently manifests with visual field defects in the early stage only at the peripheral retina, gradually progressing towards the central visual field while diabetic retinopathy and macular degeneration often impact the foveal region initially. Therefore, measurement of localized inner retina metabolic change with focal oximetry in smaller vessels, even down to the capillary level, is highly desirable for better disease characterization.

Therefore, our overall motivation is to perform human retinal oximetry in capillary level vessels less than100 µm in diameter by vis-OCT angiography (vis-OCTA). The paper consists of two major components. First, we implemented vis-OCTA in human retina to aid 3D localization of small vessels and capillaries. Given the sensitivity of our system, we optimized the scanning protocol to achieve a 3×7.8 mm^2^ field of view. Second, we performed microvascular oximetry at the macular region, focusing on individual arterioles and venules immediately above the capillary network and then the capillary network itself after spatial averaging. The measured oximetry values were validated based on the known alternating pattern of feeding arterioles and draining venules in the macula. The sO_2_ from the capillary network was found to be intermediate between measured values of the arterioles and venules.

## Methods

### System

The vis-OCT system is modified from our previous visible and near infrared OCT setup, with only the visible light channel operating (Fig. 1a) [7]. Two edge filters selected the illumination wavelength range from 545 to 580 nm (Fig. 1b, 1c). We intentionally used a narrower bandwidth than our previous design to increase the power spectral density for subsequent vis-OCTA processing. Within the bandwidth, the large spectral contrast between oxy- and deoxy-hemoglobin can still be captured (Fig. 1b). The sample arm consisted of a fiber collimator (*f* = 7.5 mm), a pair of galvanometer mirrors (GVS002, Thorlabs, USA), and a 2:1 telescope relay system. We also installed an electronically tunable lens (EL-3-10, Optotune AG, Switzerland) for a fast-focusing adjustment. We used a custom-made spectrometer to record the spectral interferograms. The spectrometer consists of a fiber collimator (f= 60mm), a 2400 lines/mm transmission diffraction grating (Wasatch Photonics, USA), a focusing lens (f = 150mm), and a high speed line scan camera (E2V Octoplus, Teledyne, France). The laser power on the pupil was less than 0.25mW, and the A-line speed is 100 kHz with 9.7 µs exposure time for each A-line. We characterized the roll-off performance of the system using a mirror reflectance as the sample, and an ND filter (OD=2) to attenuate the sample signal (Fig. 1d). The sensitivity is estimated to be ∼82dB and the roll-off speed is ∼6 dB/mm in air. The axial resolution is measured to be ∼5 µm close to the zero-delay line.

**Fig. 1.**
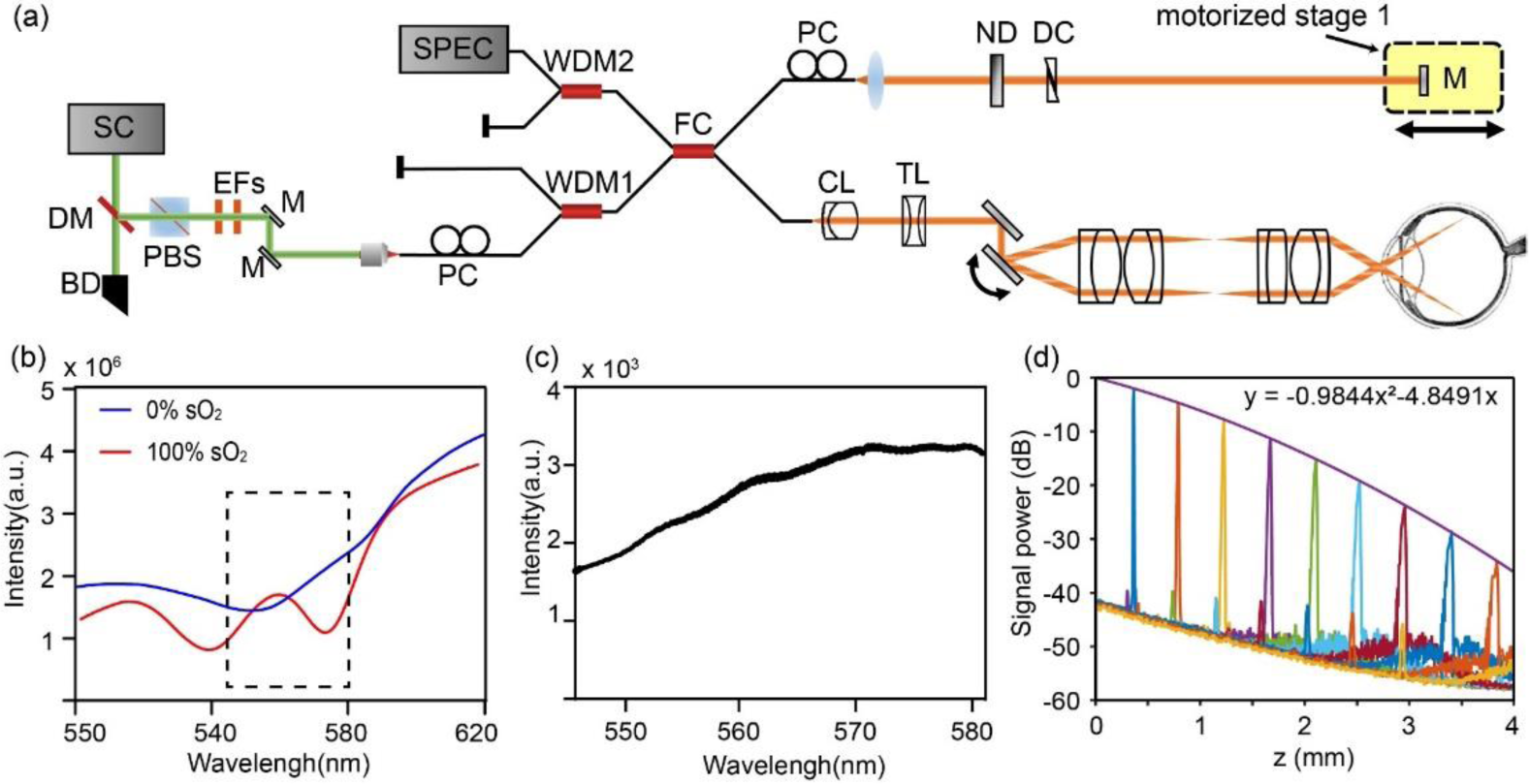
The system characteristics. (a) Schematic of the vis-OCTA system. SC: supercontinuum source; DM: dichroic mirror; BD: beam dump; PBS: polarization beam splitter; EF: edge filter; M: mirror; PC: polarization controller; WDM: wavelength division and multiplexer; FC: fiber coupler; CL: collimating lens; TL: tunable lens; ND: neutral density filter; DC: dispersion controller. (b) The wavelength selection of the spectrometer. Simulated scattering signal from the oxygenated and deoxygenated blood. (c) The measured light source spectrum. (d) System roll off characterization.

All the experimental procedures were approved by the Boston Medical Center Institutional Review Board and adhered to the tenets of the Declaration of Helsinki. Informed consent was obtained prior to imaging.

### Vis-OCTA imaging and processing

The scanning protocol is conventional and modified from our previous paper [28]. The fast scanning galvanometer mirror was controlled by an 80% duty cycle triangle wave. The slow scanning was controlled by a ramping wave. At each slow scanning location, repetitive frames were taken to harness the motion contrast of blood flow for vis-OCTA. There are three essential parameters in the imaging protocol: numbers of repetitive frames (*N*_*rep*_), the time interval between two adjacent frames (Δ*T*) for the motion contrast, and the A-line scanning density on the retina (*Σ*_*Aline*_, μm^-1^). These three parameters altogether determine the vis-OCTA ima0067e quality and the field of view (FOV), and their optimization is discussed in the Results section.

The method to generate wavelength-dependent four-dimensional(*x, y, z, λ*) vis-OCTA and vis-OCT has been described before [25,29]. We combined the split-spectrum OCTA and the complex-signal-based optical microangiography to enhance the motion contrast [30,31]. The split-spectrum method also provides the spectral information for sO_2_ calculation.

The original raw spectrograms were pre-processed by normalizing the source spectrum, a DC removal, *k*-space resampling, and a digital dispersion compensation for improving the image sharpness [32]. A Gaussian spectral window in the wavenumber *k* domain (FWHM *k*= 0.11 μm^-1^) was then used to sweep the whole interferogram in 11 steps to generate the wavelength-dependent complex OCT signal after a series of Fourier transforms. The differential contrast between two adjacent frames at one slow scanning location was taken based on the complex OCT signal, and then the absolute value was taken as the OCTA signal. When *N*_*rep*_ is larger than 2, all the OCTA signals from two adjacent B-scans were averaged to ensemble an OCTA frame

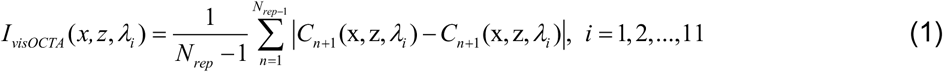

where *C*(*x, z, λ*_*i*_) is the complex B-scan image at each spectral window *λ*_*i*_. After vis-OCTA were generated for each Gaussian spectral window, we averaged them to compile one 3D vis-OCTA image.

For wavelength-dependent vis-OCT structural images, we simply average the absolute values of *N*_*rep*_ frames at each B-scan location, instead of taking differential contrast as in OCTA.

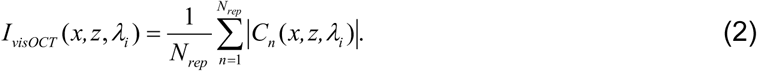

### Data processing for oximetry in small vessels

We used the following workflow to extract the microvascular spectral signal and calculate the sO_2_ (Fig. 2a). The first step is to generate wavelength-dependent vis-OCT and vis-OCTA four dimensional (*x, y, z, λ*_*i*_) datasets (Fig. 2b), as described above. The second step is to locate the blood vessels through *en face* and axial segmentations. The third step is spectral normalization and extraction to estimate sO_2_.

**Fig. 2.**
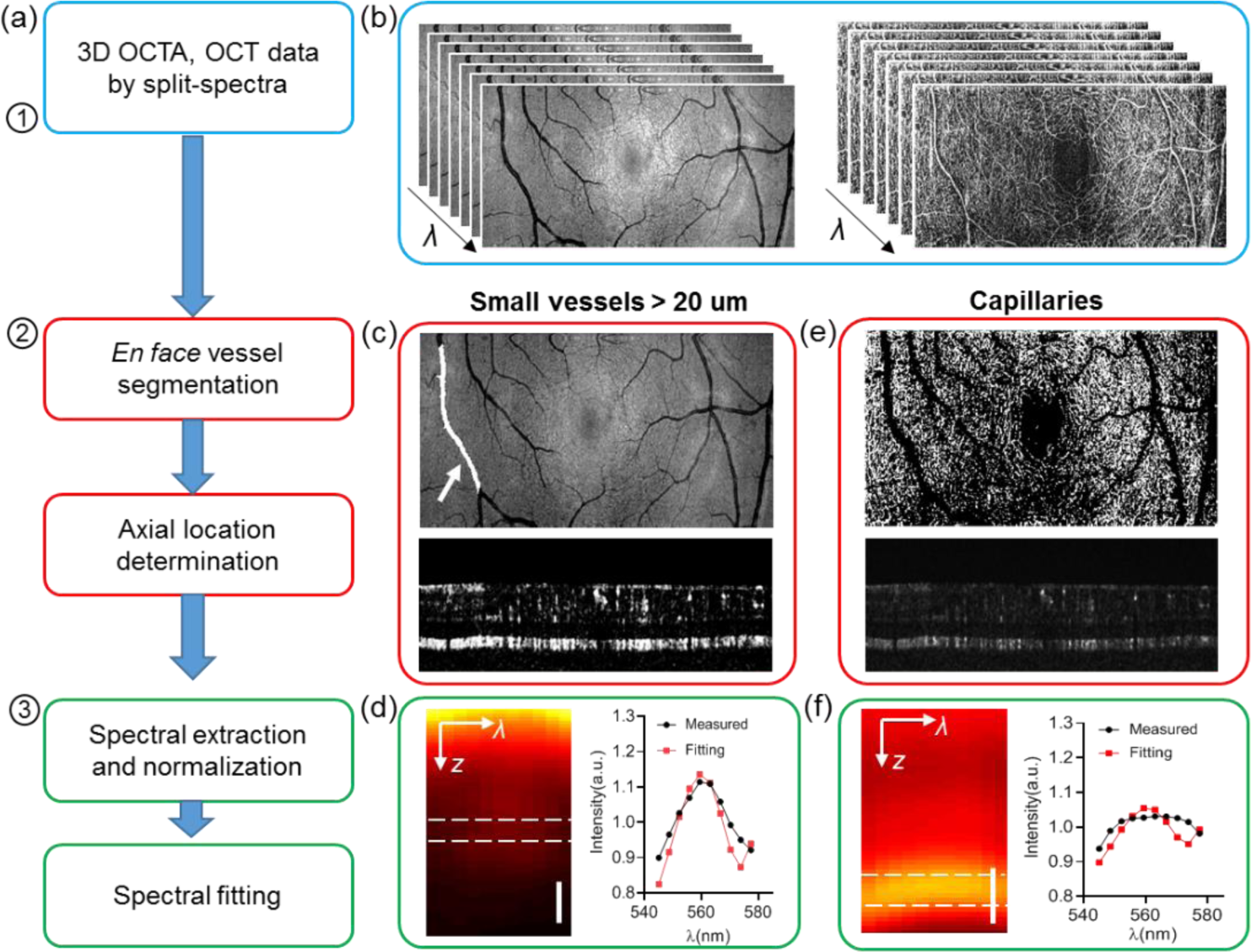
Data processing flow for small vessel and capillary sO_2_ calculation. (a) Flow chart of the data processing. (b) Wavelength-dependent vis-OCT and vis-OCTA generated by a split spectra method. (c) Illustration of the vessel segmentation on the lateral projection image from vis-OCT, and vessel bottom selection on vis-OCTA for small vessels > 20 μm. (d) An example of the averaged spectrograph in terms of depth within a vessel, and the spectrum extracted from the vessel bottom for the spectral fitting (red). (e) Illustration of capillary segmentation from *en face* vis-OCTA projection and the depth locations. (f) An example of a spectrogram on capillaries, and the spectrum extracted from the spectral fitting.

#### Vessel segmentation

For individual arterioles and venules, the *en face* maximum intensity projection (MIP) of vis-OCT was used to locate the lateral ROI for the vessels. The vessel ROI was manually segmented and stored as the vessel mask (Fig. 2c). The axial location of the vessel bottom was identified in the vis-OCTA B-scans. We detected the retinal surface by setting an intensity threshold from the structural vis-OCT B-scans. The same surface was used in both vis-OCT and vis-OCTA datasets. We then flattened the retina surface, and located the vessel bottom by the lower boundary of the vis-OCTA signal within each vessel segment (Fig. 2c). For each vessel segment, a constant depth selection was used for the spectral extraction assuming a consistent vessel size.

the capillary flow signal from the *en face* MIP of vis-OCTA was used to isolate the capillary network. The larger vessels were marked out as shown in Fig. 2e. Then we binarized the image to serve as the capillary ROI. To locate the depth of the capillaries, the axial position of the maximum value along each vis-OCTA A-line within the capillary ROI was recorded.

#### Spectral extraction and normalization

After 3D segmentation, a spectrogram in terms of depth was generated. For individual arterioles and venules, the vessel surface was flattened and all vis-OCT A-lines within the vessel lateral ROI were averaged to generate a spectrogram in *z* and *λ* (Fig. 2d). Then the spectrum was obtained by averaging signals from a depth range of 10μm centered at the vessel bottom. To remove systemic bias due to propagation through the eye, we further performed a spectral normalization by the spectrum from the non-vascular nerve fiber layer tissues. For capillaries, we flattened all the capillaries and averaged the signal within the capillary lateral ROI to generate the spectrogram in *z* and *λ* as shown in Fig. 2f. The signals from a depth range of 10μm centered on the capillaries were averaged to extract the spectrum, which was further normalized by the spectrum averaged from the inner plexiform layer and the inner nuclear layer excluding the capillaries.

#### Spectral fitting to estimate sO_2_

After the spectra were extracted and normalized, we used a previously published method to calculate sO_2_ by matching the experimental spectra with the model [4,25]:

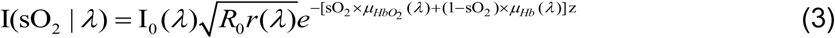

in which *I*_*0*_*(λ)* is the spectrum of the light source; *R*_0_ is the reflectance of the reference arm and assumed to be a constant; and *r*(λ) (dimensionless) is the reflectance at the vessel wall, the scattering spectrum of which can be modelled by a power law *r*(*λ*) = *Aλ*^*-α*^, with *A* being a dimensionless constant and *α* modelling the decaying scattering spectrum from the vessel wall. The optical attenuation coefficient *μ* (mm^-1^) combines the absorption (*μ*_*a*_) and scattering coefficients (*μ*_*s*_) of the whole blood, which is both wavelength- and sO_2_-dependent:

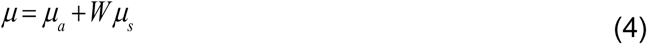

where *W* is a scaling factor for the scattering coefficient at *W* = 0.2. Hb and HbO_2_ denote the contribution from the deoxygenated and oxygenated blood, respectively. *z* denotes the light-penetration depth through vessels. When taking a logarithmic operation, Equ 2 turns to a linear equation to *μ*,

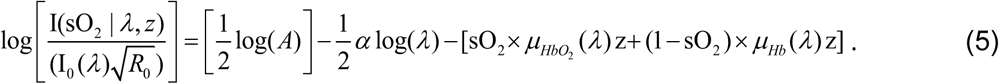

Then a least square fitting was performed to match experimental and model-based spectra to estimate sO_2_.

## Results

### Imaging protocol optimization

We first evaluate the effect of *N*_*rep*_ on vis-OCTA’s signal-to-noise ratio (SNR), defined by

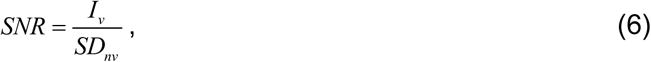

where *I*_*v*_ is the vis-OCTA intensity on blood vessels, and *SD*_*nv*_ is the image intensity standard deviation within the background non-vascular areas [33]. We collected a dataset by *N*_*rep*_ = 12, and processed vis-OCTA images from the first *N*_*rep*_ frames for *N*_*rep*_ = 2, 4, 6, 8, 12. For example, the case of *N*_*rep*_ = 2 only processes the first two frames and so on. This approach ensures that the other experimental conditions are virtually identical. We can see the image quality improvement when averaging more vis-OCTA frames (Fig. 3a-3e). The image SNR increases more rapidly in the beginning and slows down when *N*_*rep*_ *> 6* (Fig. 3f). While the image quality improves with increasing *N*_*rep*_, the acquisition time proportionally increases as well. Therefore, we chose *N*_*rep*_ *= 3* for the final imaging protocol to balance the image SNR and the total scanning time. The image is 320 by 128 pixels.

**Fig. 3.**
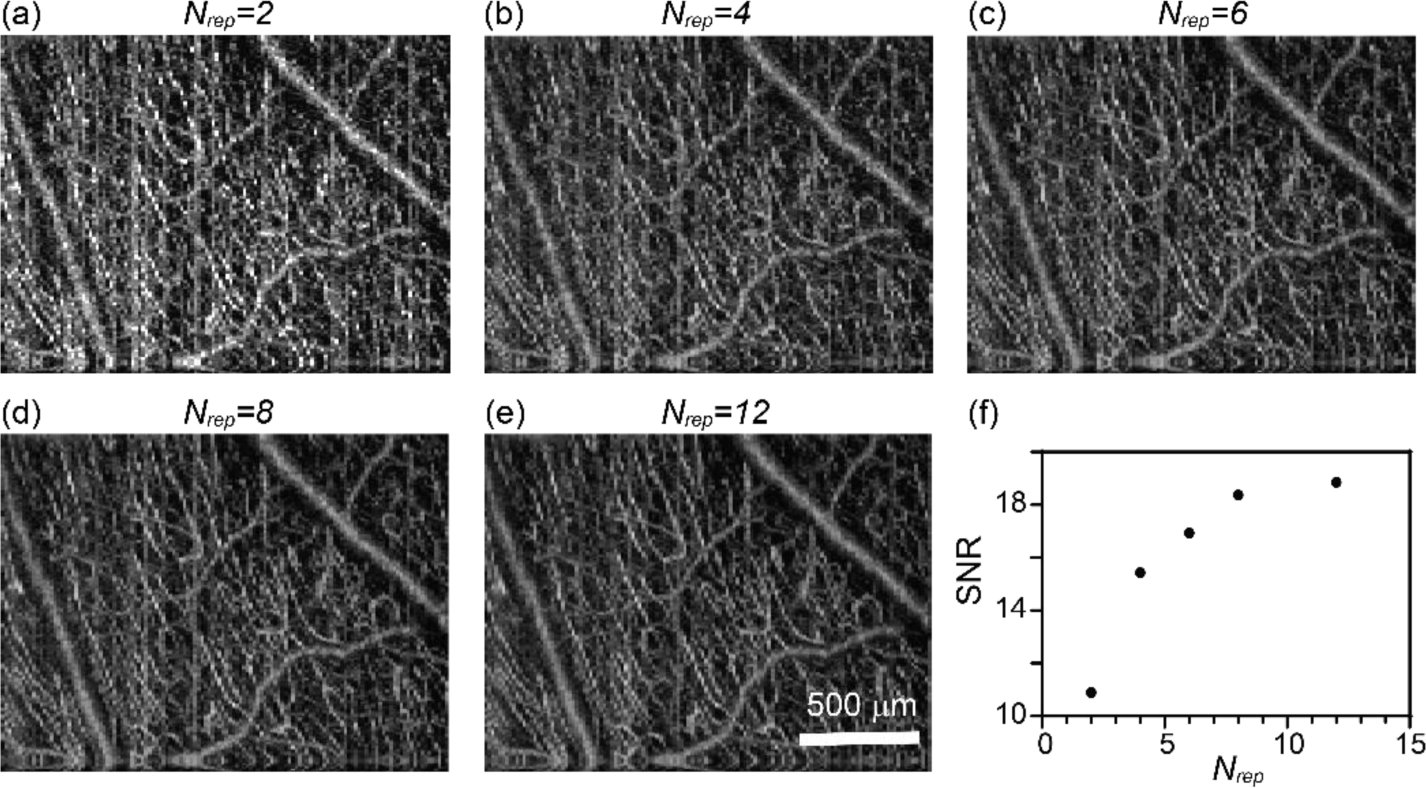
The effect of the repetition times on human retina vis-OCTA. (a-e) En face maximum intensity projection of vis-OCTA in the inner retina at repetition times *N*_*rep*_ equal to 2-12 at each B-scan location. (f) Capillary vis-OCTA SNR versus *N*_*rep*_.

We next evaluated the effect of ΔT on the image quality. The A-line scanning density was kept constant at 0.13 μm-1, while ΔT increased from 1 ms to 7 ms by increasing the number of A-lines per frames from 80 to 560. As a result, the length in the fast scanning direction increases as well (Fig. 4a-4d) from 0.6 to 4.3 mm, respectively. The same FOV was imaged in all cases in Fig. 4a, so that the SNR calculation could be repeated at the same imaging location. We noticed that the image contrast changes non-monotonically, increasing until ΔT = 5ms and decreasing at ΔT = 7 ms. The initial SNR increase may be due to the larger decorrelation between two repeated frames when ΔT increases. The contrast then deteriorates when ΔT becomes longer than 5ms as clear motion artifacts start to appear (*e.g.* vertical stripes in Fig. 4c, d). To balance the FOV and the image quality, we chose ΔT = 5ms for our imaging protocol. At 100 kHz A-line rate and 80% duty cycle of the fast scanning, ΔT = 5ms determines the number of A-line per B-scan in our protocol to be 400, resulting in a ∼3mm scanning range in fast axis.

**Fig. 4.**
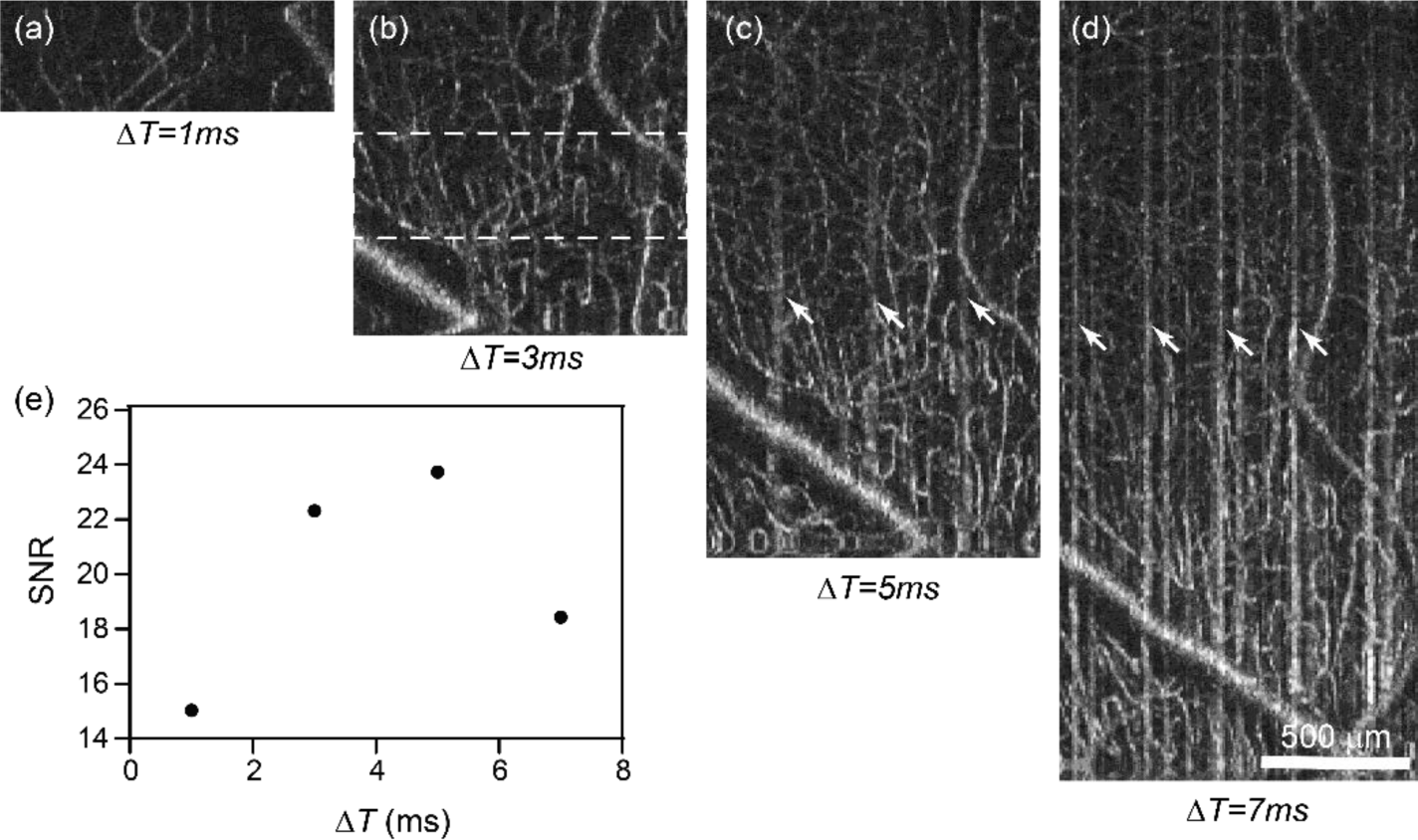
The effect of the B-scan interval time on human retina vis-OCTA. (a-d) *En face* maximum intensity projection of vis-OCTA in the inner retina at the interval time *ΔT* equal to 1-7 ms between two consecutive frames at each B-scan location, while the A-scan density were maintained constant. (e) Capillary vis-OCTA SNR versus *ΔT.*

Lastly, we evaluated the effect of A-line scanning density on the vis-OCTA SNR. We kept *N*_*rep*_ and *ΔT* constant at *N*_*rep*_ = 3 and *ΔT = 4 ms* with 320 A-line per B-scan. The length of the scanning was controlled by the amplitude of the triangle wave fed to the fast scanning galvanometer mirror. Based on the scanning angle at the pupil entrance and the approximated eyeball diameter of ∼25mm, we varied the scanning length from approximately 0.375 to 3.375 mm in five acquisitions, corresponding to the A-line scanning density *Σ*_*A-line*_ = 0.85, 0.29, 0.17, 0.12 and 0.09 μm^-1^. As Fig. 5 shows, the image SNR decreases with the decreased A-line scanning density. This effect is likely due to the fringe washout effect as each exposure integrate signals from the scanning length per pixel, and becomes more prominent as the scanning length increases. Then, we fixed *Σ*_*A-line*_ *=* 0.13 μm^-1^, ΔT = 5 ms, and 400 A-lines per frame for the final imaging protocol, which yielded an imaging length on the fast scanning direction of ∼ 3 mm.

**Fig. 5.**
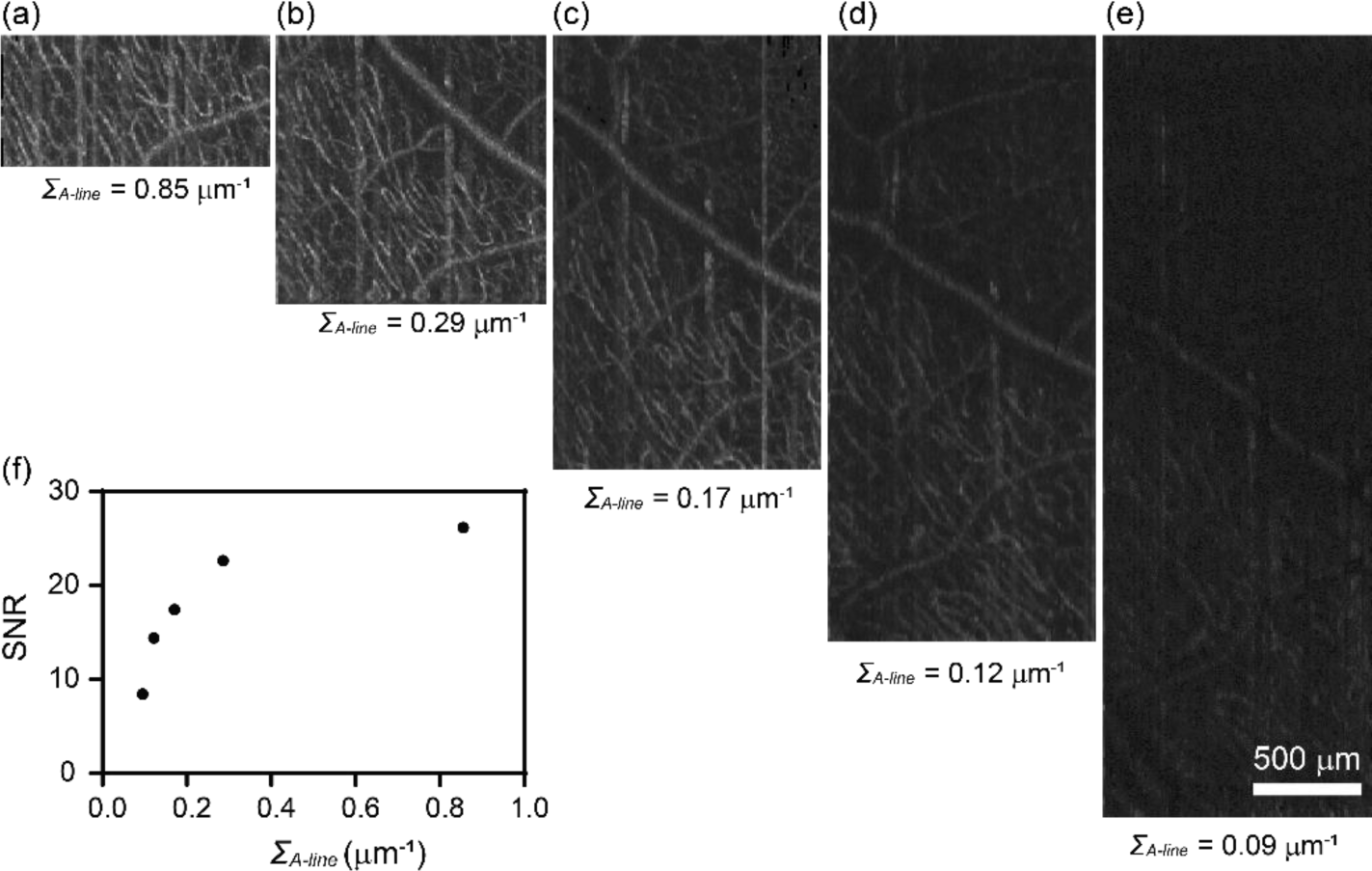
The effect of the A-line scanning density within a B-scan on human retina vis-OCTA. (a-e) *En face* maximum intensity projection of vis-OCTA in the inner retina at the A-line scanning density *Σ*_*A-line*_ equal to 0.85-0.09 μm within a B-scan, while *N*_*rep*_ and *ΔT* were maintained constant. (f) Capillary vis-OCTA SNR versus *Σ*_*A-line*_.

### Human retina vis-OCTA

After optimizing the protocol, we performed vis-OCTA on a human volunteer as shown in Fig. 6. The FOV at one single acquisition takes 400 x 400 pixels to cover ∼3×7.8 mm in the retina. The *N*_*rep*_ = 3 and the total time for one acquisition is 6 s. Figure 6a shows a wide field vis-OCTA mosaic including 11 acquisitions. Detailed microvasculature down to individual capillary-level can be visualized in two magnified images (Fig. 6b, 6c). After the split-spectra processing, the depth resolution was relaxed to ∼27 μm, sufficient to isolate the superficial, intermediate, and deep capillary plexuses at the macula (Fig. 1 in Appendix). The capillary contrast is higher within the nerve fiber arcade, which is consistent with the high metabolic demand in the retinal nerve fiber layer and the ganglion cells.

**Fig. 6.**
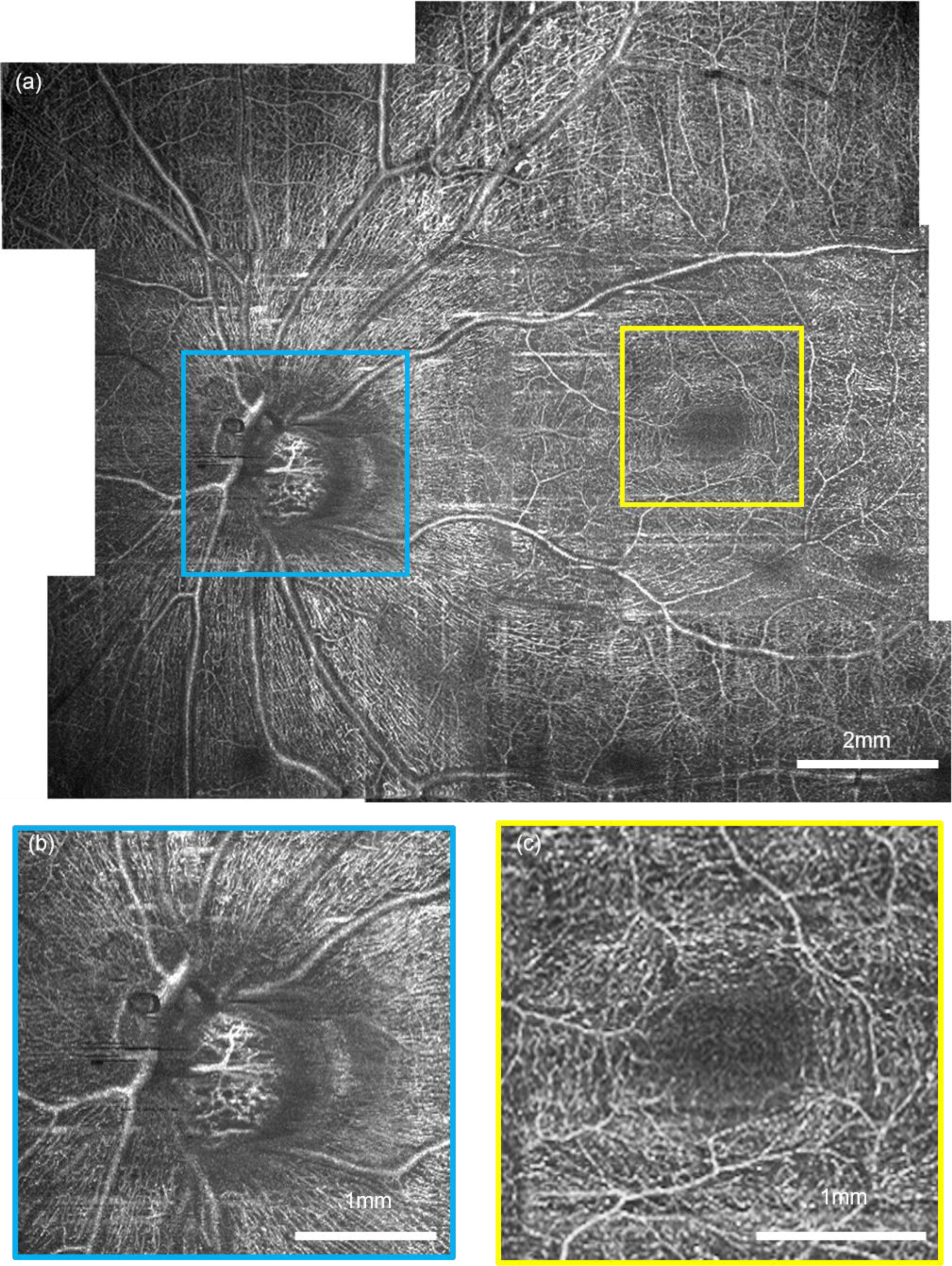
Human retina vis-OCTA. (a) A mosaic wide-field vis-OCTA *en face* projection made of 10 acquisition on a healthy volunteer aged 37. (b-c) The magnified details of the capillary network at the optic nerve head and fovea.

### Microvascular sO_2_ in human retina in vivo

We performed vis-OCT microvascular oximetry at the foveal region as demonstrated in Fig. 7. We color-coded the vis-OCTA images with sO_2_ values for small vessel segments and the whole capillary bed (Fig. 7a). The alternating pattern of feeding arterioles and draining venules is apparent around fovea as demonstrated by the sO_2_ estimations. As expected, the averaged sO_2_ value over the capillary bed is intermediate between the values of the arterioles and venules. We plotted a representative spectrogram from an arterial segment, a venule segment, and the underlying capillary network (Fig. 7b-7d). Figures 7b and 7c have a high intensity at the vessel surface that attenuates when penetrating through the vessels. The depth at the vessel bottom provides sufficient signal contrast to extract reliable spectra. Figure 7d exhibits a different pattern with the strongest signal appearing within the capillaries as expected based on the stronger capillary contrast in structural vis-OCT images [34,35].

**Fig. 7.**
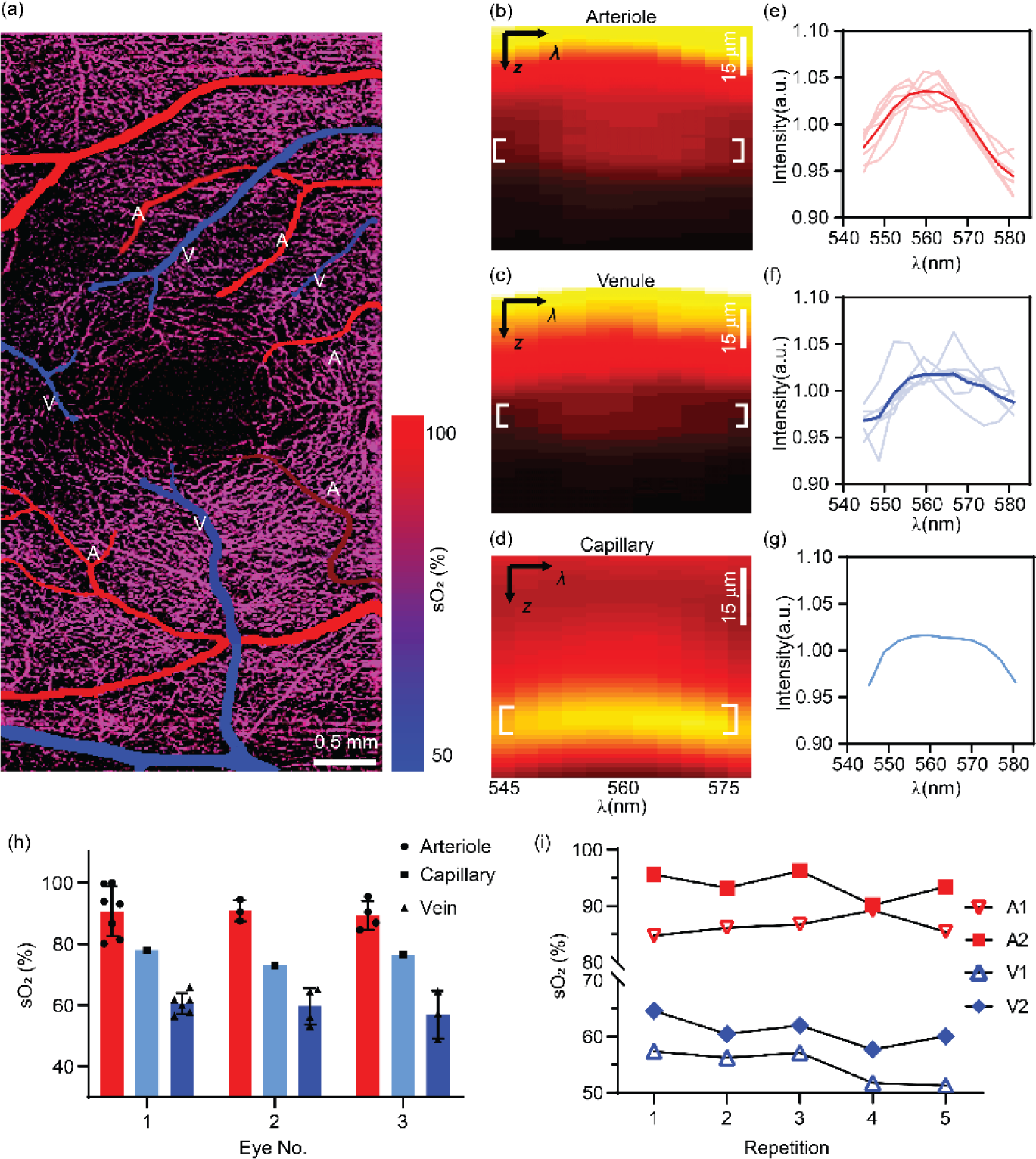
Human retinal oximetry on small vessels and capillaries by vis-OCTA. (a) Color-coded by sO_2_ on arterioles, venules and capillaries at the perifoveal region. (b-d) Representative spectrogram from an arteriole, a venule and the capillary network from panel (a). (e-f) The measured spectra from all arterioles and venules, and capillary network. The solid and light curves were the averaged and individual spectra from all the vessels in panel (a). The capillary spectrum were averaged within the entire field-of-view as shown in panel (a). (h) sO_2_ calculations from three healthy eyes. Bar = std. (i) The longitudinal repeatability from different vessels from a same healthy eye.

In Fig. 7e-7g, we plotted the spectra from all the arterioles, venules, and capillary beds displayed in Fig. 7a. The spectral contrast of the blood resembles that seen with the oxygenated blood plotted in Fig. 1b, showing a peak around 560 nm. The value of sO_2_ estimation by spectral fitting is then plotted in Fig. 7h as the 1^st^ subject. We further performed the same method on two additional healthy eyes, and their results are also summarized in the same graph, and Table 1. Overall, the mean values of arterial, venous, and capillary sO_2_ are quite consistent among three different subjects.

**Table 1.**
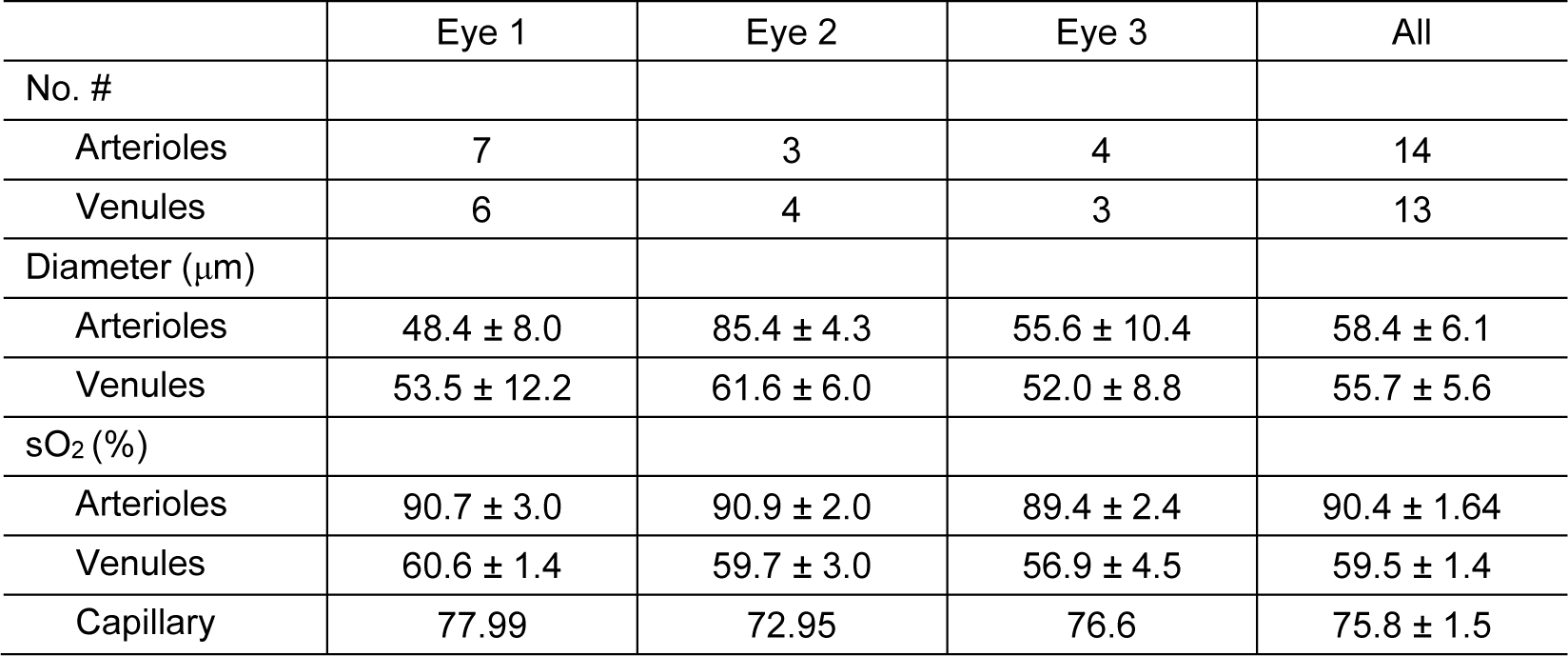
Summary of microvascular sO_2_ measurements at the perifoveal region from three healthy eyes

We also tested the repeatability of the microvascular sO_2_ by vis-OCTA (Fig. 7i). A total of 5 measurements were taken from the same eye on three different dates. The first three repetitions are taken on the same day, and the other two are taken on different days spanning over 5 weeks. The coefficients of variation (CV) for all measured vessels are < 3% within the same day and < 5% over all the repetitions.

## Discussion

In this paper, we describe the first vis-OCTA imaging of human retina using a redesigned spectrometer and optimized imaging protocol allowing for enhanced vascular signal and localization of the microvasculature. Using our methods, we are capable of measuring sO_2_ in vessels smaller than 100 μm, all the way down to the capillary network. We demonstrate the robustness and repeatability o sO_2_ calculation. The unique advantage of vis-OCTA oximetry is the capability of excluding confounding factors from other retinal layers through 3D segmentation. In addition, the accurate vessel localization of vis-OCTA allows for measurement of capillary sO_2_, which is unattainable by fundus-based oximetry.

The limitation of this study is the lack of independent validation of our sO_2_ calculation. In our previous studies on rodent retinal oximetry, a pulse oximetry was used to serve as a gold standard for arterial sO_2_ close to the ONH [25,29]. sO_2_ alterations due to oxygen diffusion as blood travels through the retina would limit this approach for validation of measurements in microvasculature distant from the ONH. From our calculations, we observe the clear separation of alternating feeding arterioles and draining venules, which serves as a physiologic and anatomic validation. Yet we still lack a gold standard for capillary sO_2_. Further physiologic (*e.g.* hypoxia challenge) or pathologic validations with known impact on oxygen levels are warranted to rigorously validate the accuracy and sensitivity of microvascular sO_2_ by vis-OCTA.

We should also emphasize that all the spectral extraction was performed on structural vis-OCT signals with vis-OCTA used to provide vessel localization. Because of the stringent requirements for imaging speed and SNR for vis-OCTA, the FOV is limited to 3×7.8 mm^2^ at each acquisition with 6s total acquisition time at the current system characterization. This may be a limiting factor for large-scale clinical applications. Alternatively, we could use a dual-channel system with both visible and conventional near infrared (NIR) wavelengths to simultaneously acquire OCTA images [7,36]. While the NIR channel could provide rapid OCTA with a larger FOV for vessel localization, the co-registered vis-OCT structural data could provide spectral contrast for sO_2_.

To summarize, we have demonstrated first visible light optical coherence tomography angiography for human retinal imaging. By achieving a 100 kHz A-line rate, the fastest acquisition speed reported so far for human retinal vis-OCT, we can acquire a field of view (FOV) of 3×7.8 mm^2^ image within 6 seconds. The angiography also enables accurate localization of microvasculature to allow us to perform human retinal oximetry in vessels less than100 µm in diameter. This technical advance lays a foundation for assessing human retinal local oxygen extraction and metabolism.

## Appendix

**Table 2.**
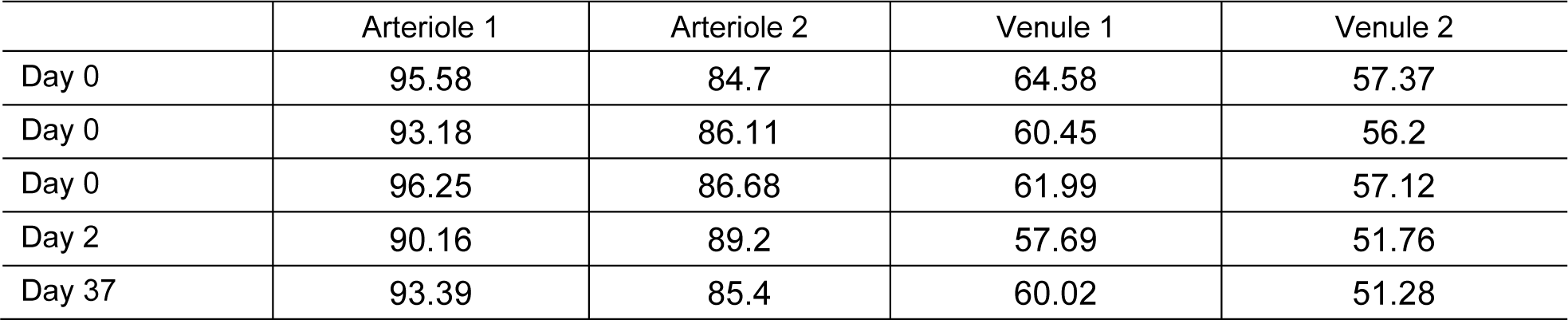
The repeatability of the microvascular sO_2_ (%)

**Fig. 1.**
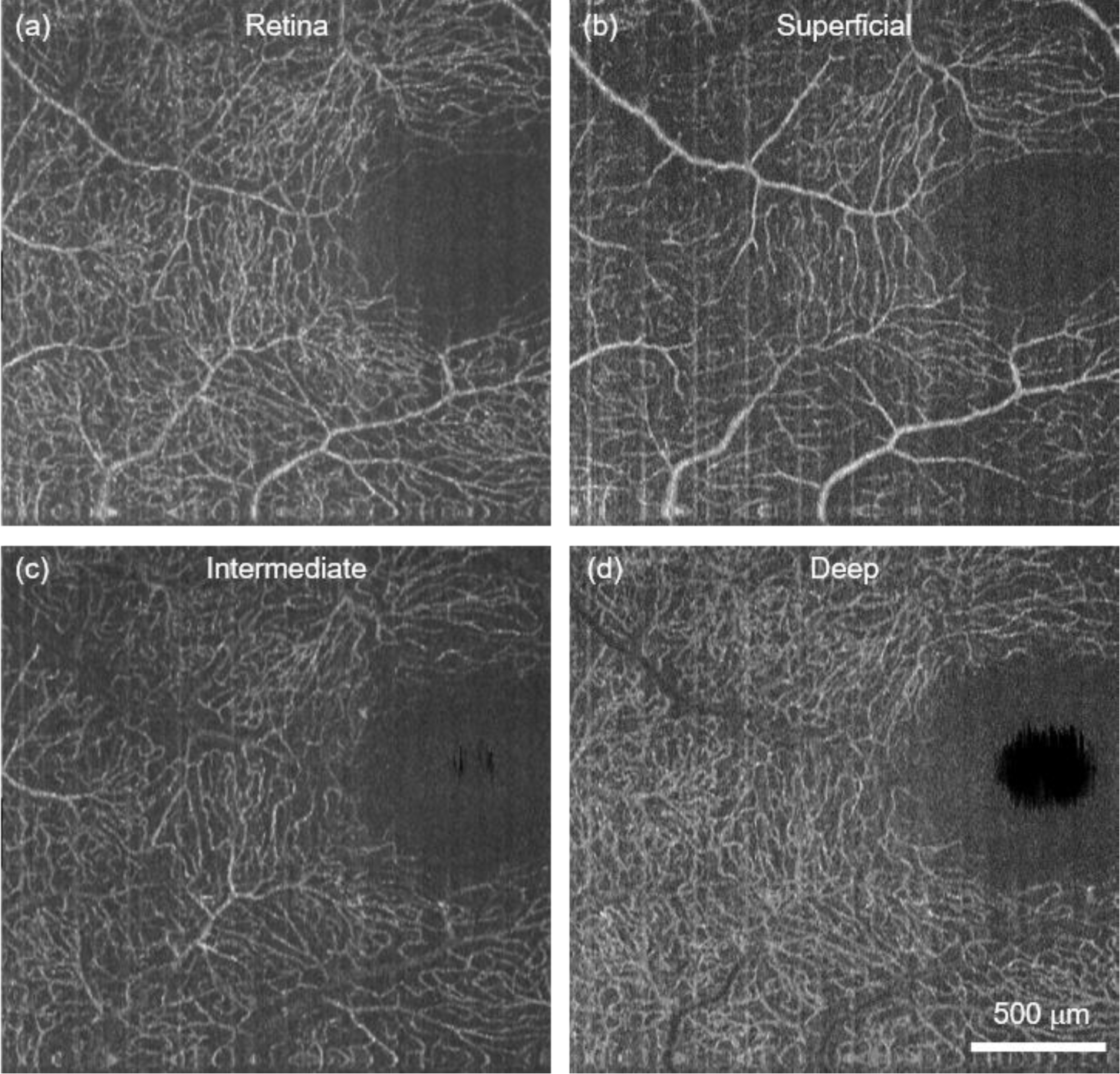
(a) Human retinal capillaries at macula by vis-OCTA. (b-d) The superficial, intermediate, and deep capillary plexus isolated from (a).

